# Ghost lineages can invalidate or even reverse findings regarding gene flow

**DOI:** 10.1101/2022.01.06.475184

**Authors:** Théo Tricou, Eric Tannier, Damien M. de Vienne

## Abstract

Introgression, endosymbiosis and gene transfer, *i*.*e*. Horizontal Gene Flow (HGF), are primordial sources of innovation in all domains of life. Our knowledge on HGF relies on detection methods that exploit some of its signatures left on extant genomes. One of them is the effect of HGF on branch lengths of constructed phylogenies. This signature has been formalized in statistical tests for HGF detection, and used for example to detect massive adaptive gene flows in malaria vectors or to order evolutionary events involved in eukaryogenesis. However these studies rely on the assumption that ghost lineages (all unsampled extant and extinct taxa) have little influence. We demonstrate here with simulations and data re-analysis, that when considering the more realistic condition that unsampled taxa are legion compared to sampled ones, the conclusion of these studies become unfounded or even reversed. This illustrates the necessity to recognize the existence of ghosts in evolutionary studies.

## Introduction

Gene flow between taxa affects all domains of Life, and occurs between taxa separated by different evolutionary time-scales: from introgression between populations to trans-phylum endosymbiosis. Introgression, where gene flow results from hybridization followed by repeated back-crossing with one of the parent species, is recognized as a major source of genetic variation in natural populations, that contributed to adaptation and adaptive radiation in most plant and animal groups, including humans [see 1 for a recent review]. Horizontal Gene Transfer (HGT), where gene flow occurs between distinct taxon in a non-vertical manner and is mediated by mechanisms as diverse as natural transformation, transduction, conjugation, recombination or endosymbiosis [2,3], appears now as a primary source of innovation in most taxonomic groups. Emblematic cases go from the acquisition of antibiotic resistance genes in bacteria [4], to the emergence of mitochondria and chloroplasts in eukaryotic cells by endosymbiosis [5]. These events, hereafter referred to as horizontal gene flow (HGF) can be detected with phylogenetic approaches. Signature of HGF may be seen when phylogenies reconstructed from portions of the genome that were horizontally acquired (from a few nucleotides to full chromosomes) contradict the evolutionary history of the taxa analyzed (represented by the species tree), and/or show differences in branch lengths. Indeed, the genetic (and thus phylogenetic) distance between the lineages involved in the flow (donor and recipient) for the horizontally acquired sequence reflect the time since the flow, not the time since their speciation. These simple expectations form the basis of numerous studies of HGF, either to propose new methods for the detection of gene flow [6–12], or to resolve evolutionary puzzles in various taxonomic groups [12–16].

Here, we focus specifically on the link between HGF and phylogenetic tree branch lengths. We show that the way branch lengths are interpreted in several important studies of gene flow overlooks the possible impact of ghost lineages on the expectations given above. By ghost lineages, we mean any taxon that is absent from the analysis, but is susceptible to interfere with the analyzed taxa via HGF. This means the extant taxa that are known but not sampled, extant taxa that are unknown, and all extinct taxa. More than 99.9% of all species that have ever lived are now extinct [17], and the number of extant species that are still uncatalogued is almost an order of magnitude higher than those that have been reported (around 1.3 out of 8.7 Million eukaryotes estimated in 2011, Mora et al. 2011), or many orders of magnitudes higher according to some predictions [19]. The amount of ghost taxa in any analysis of HGF is thus potentially huge.

We reanalyzed recent studies that used the expected link between HGF and branch lengths in different contexts: for the detection of introgressed loci (the D_3_ method, Hahn and Hibbins 2019), for resolving the correct branching order of *Anopheles* species [13] and for ordering the acquisitions of genes associated with the emergence of eukaryotic cells [14,16] and the elaboration of the chloroplast membrane [20]. Using simulations involving species trees with extinctions and gene trees prone to HGF, we show that under the realistic situation where most past and present taxa are ghosts, the branch length argument in relation to HGF is misleading, and can even lead to conclusions that are the opposite of the published ones. We emphasize the need to take into account ghost lineages in evolutionary studies and propose an alternative hypothesis when using branch lengths in phylogenies to investigate HGF: any signal of HGF should be interpreted in the first instance as an indication of the presence of ghost lineages that acted as donors, and not as an indication of a gene flow between lineages present in the analysis.

## Results

### Preamble

The three families of examples we explored in this study (the three next sections) illustrate a common principle regarding the impact of ghost lineages on phylogenetic tree branch lengths after HGF: while the time where an HGF event occurs is reflected by the divergence time (or branch lengths) between the donor and the recipient species when looking at the transferred region, this relationship is lost if the donor of the transfer (or its descendants) is absent from the analysis (*i*.*e*. it is a ghost lineage). This is illustrated in Fig 1 for different scenarios of HGF involving or not ghost lineages. This has multiple consequences. First, it invalidates the expectation that HGF leads to shorter branches (compared to portions of the genome not impacted by HGF), and the absence of HGF to longer ones (compared to portions of the genome impacted by HGF). Long branches in gene trees may be the result of HGF instead of the signature of its absence (compare the no-HGF case and the scenario C in Fig 1). Second, it prevents the use of divergence times to relatively order different HGF events. Two events, early and late, may lead to short and long branches respectively, which is the opposite of what is expected in the absence of ghosts (compare the order of A and B for HGF times and for divergence times in Fig 1).

**Fig 1.**
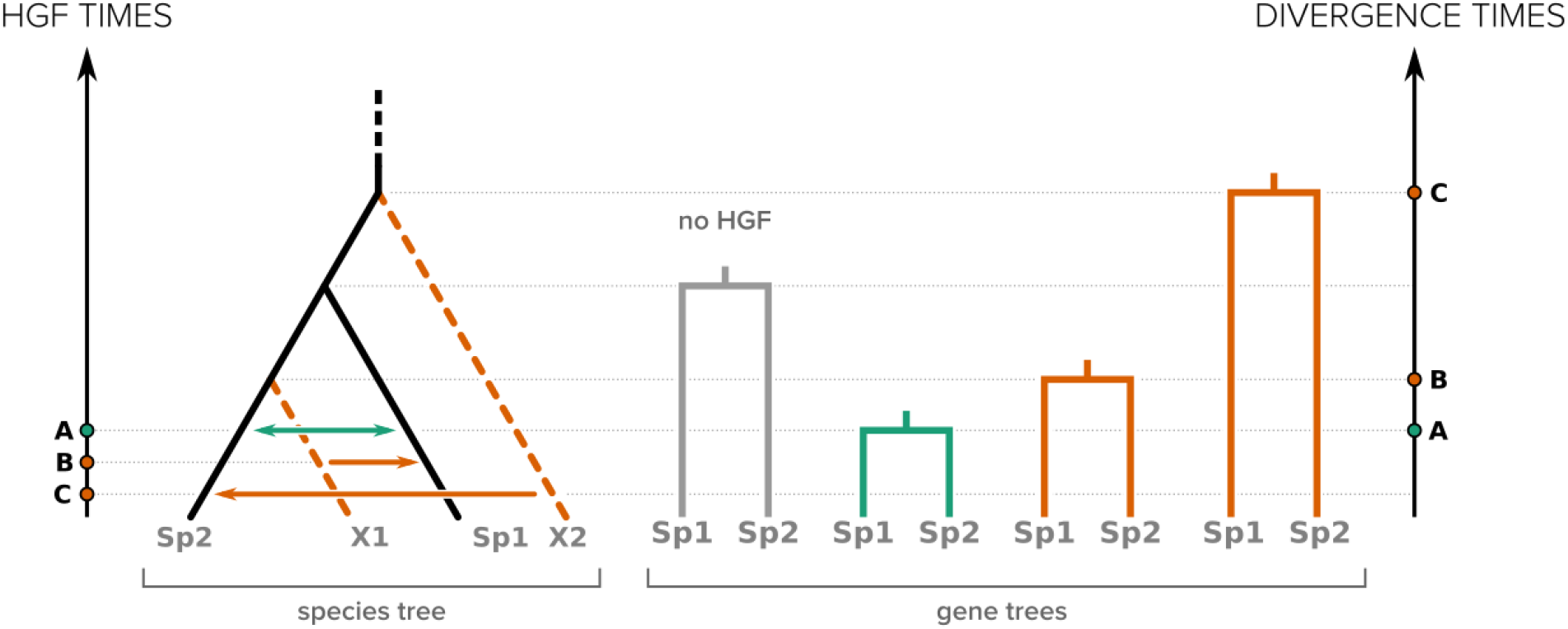
Effect of horizontal gene flow (HGF) on branch lengths in phylogenetic trees of the horizontally transferred genomic regions. Only the recipient species and its closest relative are represented in each gene tree. Green arrows depict HGF between lineages used in the analysis, while orange arrows represent HGF involving ghosts. A, B and C represent the three HGF considered.

We use simulations to explore in different contexts the impact that this loss of correlation between HGF times and branch lengths can have on studies that use branch lengths as a proxy for HGF times without considering ghosts.

### Using branch lengths to detect introgression events (the D_3_ method) is often misleading

D_3_ [9] is a recent statistical method that uses branch lengths (*i*.*e*. pairwise distances between species) to detect introgression in a three-taxon tree (Fig 2). Referring to the notations in Fig 2, the test is aimed at detecting gene flow between sp2 and sp3 or sp1 and sp3 (cases A and B, respectively), by computing the D_3_ statistics:

**Fig 2.**
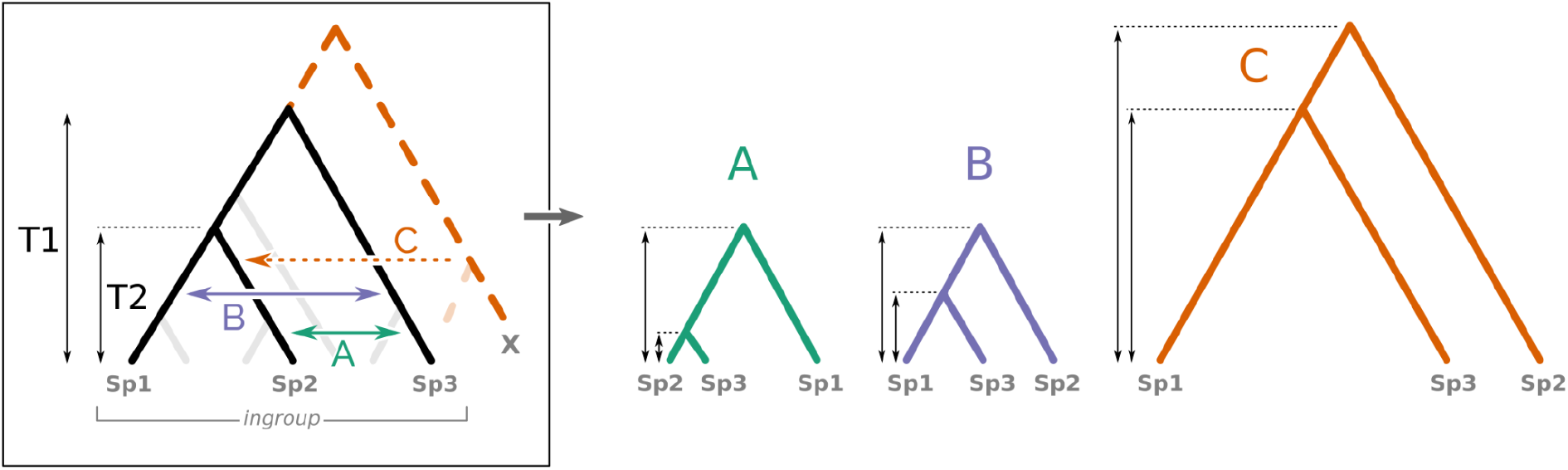
Effect of ghost lineages on a three-taxon set (sp1, sp2, sp3) as used for the D3 test of introgression. Introgression from a ghost lineage (X) from outside the ingroup of interest produces a phylogenetic tree with increased branch lengths compared to the species tree (scenario C), while branch lengths are shorter when introgression occurs within the ingroup (scenarios A and B). The new branch-lengths pattern produced by the ghost lineage (X) drastically changes the interpretation of the D3 statistical test (see text).

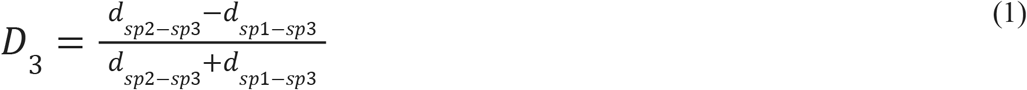

Where ***d*** represents the pairwise distance between the species, which can be obtained by a measure of genetic distance between their genome sequences (see Hahn and Hibbins, 2019 for details). According to the original description of the test, if no introgression occured, D_3_ is close to 0 regardless of the presence of Incomplete Lineage Sorting (ILS, a process where the retention of ancestral polymorphism in the populations can lead to gene trees that differ from the species tree); but in the case of gene flow, D_3_ can be either positive, revealing introgression between sp1 and sp3 (scenario B in Fig 2), or negative, revealing introgression between sp2 and sp3 (scenario A in Fig 2). The significance of the test is determined by comparison of the computed D_3_ value with a distribution of D_3_ values obtained from block-bootstrap replicates of the sequences.

Therefore the D_3_ method [9], as well as other methods based on the same principle [6,7,9–12] relies on the assumption that short branch lengths will be observed following introgression events. But what if the donor has no sampled descendants, which is probably the most frequent situation? For instance, if an introgression occurs between a taxon from outside the tree (*e*.*g*. taxon X in Fig 2) and, for example, sp2 (scenario C in Fig 2)? This introgression event increases the distance between sp2 and sp3 without affecting the distance between sp1 and sp3 and results in a positive D_3_ that is interpreted as gene flow between sp1 and sp3 even though neither is involved in the introgression. While it has been shown several times that other tests of introgression can be deceived by ghost lineages [21–23], and even though the authors of the D_3_method call for cautiousness when interpreting the results of the test because of these ghosts, the scale of this problem for the D_3_ test is still unknown.

In practice, it is possible to estimate the probability that ghost introgressions lead to erroneous interpretations of the D_3_ statistic. To this end, we simulated random species trees (40 extant species) and random introgression events with the *ms* software (see Material and Methods), assuming that the age of the root of the trees was compatible with introgressions occuring uniformly between the species in the trees, but incompatible with introgressions with species outside. This can be seen as a strict discretization of the known correlation between the probability of hybridization, and consequently of introgression, and the genetic distance between species [24–27]. We will see later why this is justified.

For all possible three-taxa samples where D_3_ was significantly different from 0, we evaluated whether this result was imputable to (i) introgression within the group containing the three taxa of interest (the ingroup), even when the donor taxon was not the one directly involved (a donor ghost taxon, sister to the one sampled, can give the same signal, see Fig 2), or (ii) a “ghost introgression” (from outside the ingroup), resulting in an erroneous interpretation of the test. Unsurprisingly, the probability of erroneously interpreting the D_3_ statistic was high when the size of the ingroup was small relative to the total size of the tree, almost reaching 100% when the ingroup comprised less than 20% of all taxa in the tree (Fig 3). Biologically speaking, it seems realistic to consider that there are more taxa with whom introgression can occur outside than inside a given three-taxa clade, especially when the three taxa chosen are “closely related species”, which is a condition for the test to be used [9].

**Fig 3.**
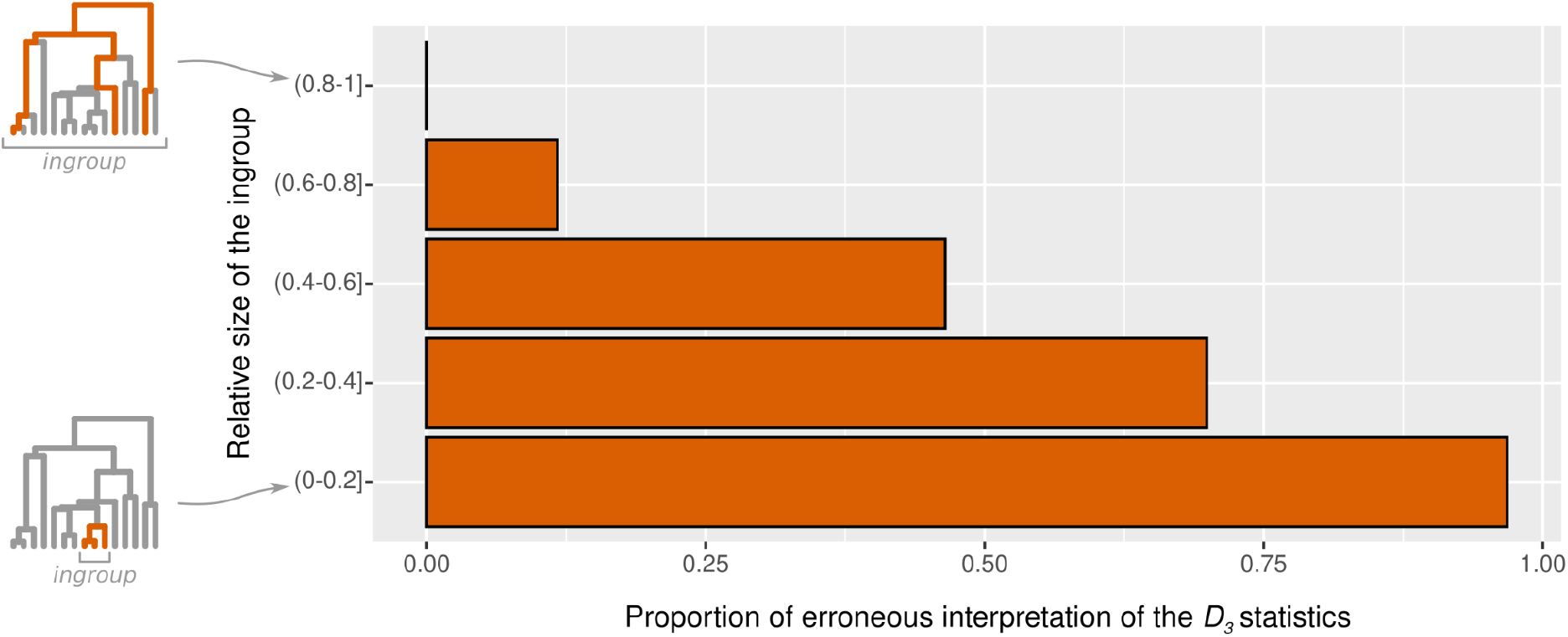
Proportion of cases (from simulations) where a D_3_ test is significantly different from 0 because of an introgression from a ghost lineage from outside the ingroup. These cases may be incorrectly interpreted (see text) and lead to the misidentification of both taxa involved in the introgression. The proportion of erroneous interpretations of the test is computed for different relative sizes of the ingroup, *i*.*e*. the number of taxa of the smallest subtree containing the three taxa studied divided by the total number of taxa in the tree (see trees on the left). Data underlying this figure can be found on Zenodo (doi: 10.5281/zenodo.6901799) and GitHub (https://github.com/theotricou/Ghost_branch_length/tree/main/1_D3).

Therefore we propose that any D_3_ statistic that is significantly different from 0 should be interpreted in the first instance as the result of an introgression event originating from outside the tree formed by the three taxa considered.

### Genes with long branch lengths are not good markers of the mosquito species phylogeny

It is common practice in phylogenetics to use gene markers that are supposed to not be horizontally transferred to construct the history of species diversification. This is important because HGF can change the branching structure (*i*.*e*. the topology) and the branch lengths of gene phylogenies (Fig 2). The assumption that HGF decreases branch lengths in gene phylogenies would suggest that among multiple possibilities of species tree topologies, the one similar to the gene trees with the longest branch lengths would represent the “true” branching order because it agrees with genes that are expected to not have experienced gene flow.

This approach, *i*.*e*. finding the species tree supported by the genes with the longest branches, was proposed and used to recover the branching order of taxa in the *Anopheles gambiae* species complex, a group of six recently diverged *Anopheles* species (Fontaine et al. 2015). The knowledge of this branching order is particularly important because it is presented, by Fontaine et al. (2015), as a prerequisite for studying introgression in this group and better understand its role in the adaptive potential of these malaria vectors. Different portions of the genome support different tree topologies, with a strong opposition between the X and the autosomal regions. To find which of these topology is the “true” species topology, the authors used the logic presented above that they detailed as follows: “Because introgression will reduce sequence divergence between the species exchanging genes, we expect that the correct species branching order revealed by gene trees constructed from non introgressed sequences will show deeper divergences than those constructed from introgressed sequences” (Fontaine et al. 2015). To perform this test, they focused on three out of the six species, *An. arabiensis, An. gambiae* and *An. melas*.

However, as explained in the preamble and in Fig 2, this logic may be incorrect because, in the presence of ghost lineages, long branches (deeper divergence) may result from introgression and are not an indication of its absence.

To test the effect of ghost lineages on the strategy proposed by Fontaine et al. [13], we simulated a species tree from which we chose three closely related species to mimic the three species of the original study that diverged only 1.8 My ago (Fontaine et al. 2015) when the age of the *Anopheles* genera is dated between 25 and 93 My [28–30]. Using *ms*, we simulated gene trees under two simple evolutionary scenarios (see Material and Methods): one involving introgression between ingroup species B and C (scenario 1), and one involving introgression from a ghost lineage from outside the ingroup to species B (scenario 2, Fig 4). We separated the simulated gene trees into two categories: those with a similar topology to the species tree and those with a discordant topology. We then computed for each category the mean divergence times (T1 and T2) for the nodes separating the three species (Fig 4). This way of presenting our results (Fig 4) mimics the Fig 3A-C of Fontaine et al. [13].

**Fig 4.**
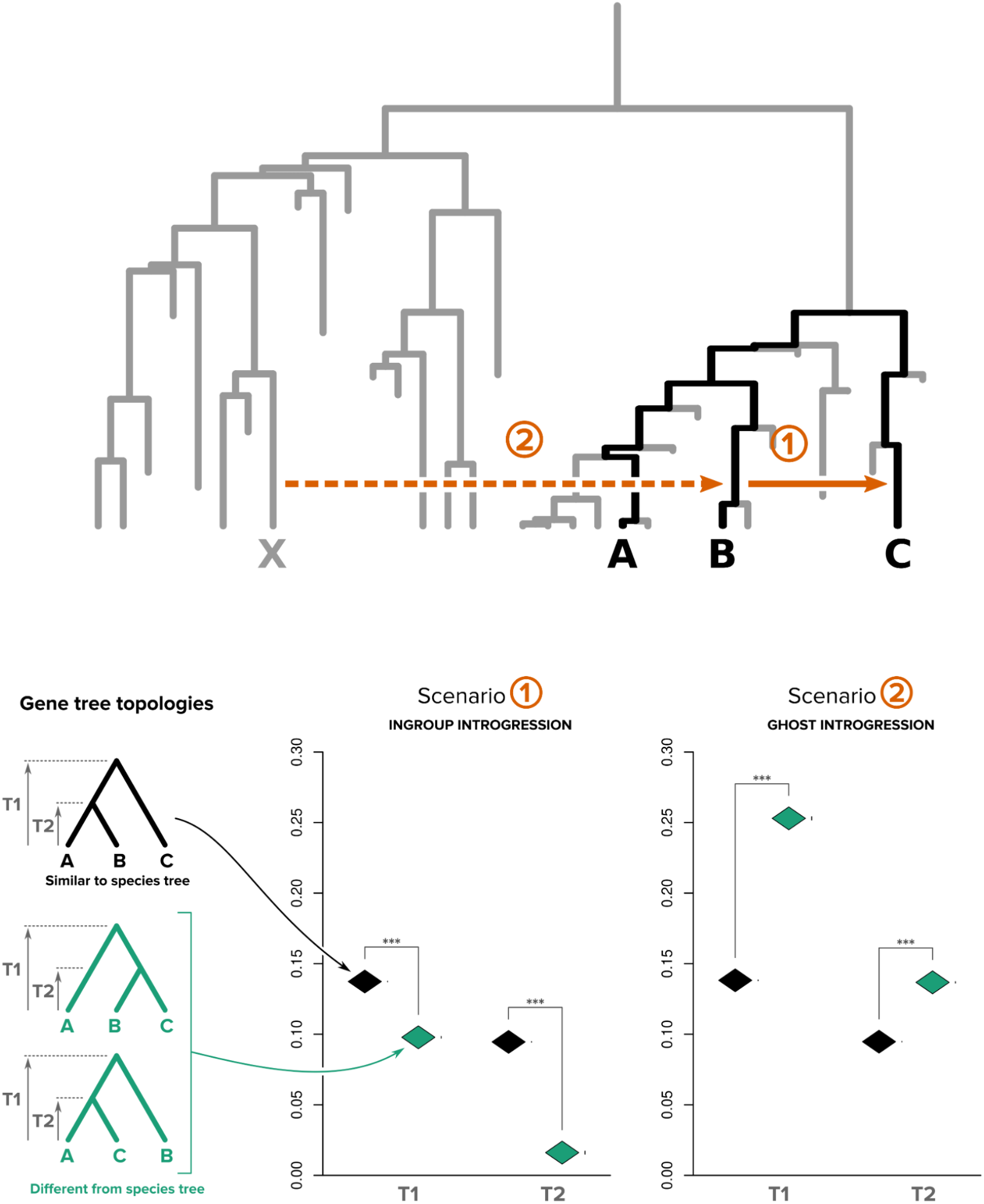
Impact of ghost lineages on the use of branch lengths to choose between alternative species trees. Top panel: the complete species tree, black branches represent the ABC tree while grey branches represent ghost lineages. Genome evolution (1,000 genes) was simulated under two different introgressions scenarios: scenario 1, ingroup introgression (solid orange arrow) between species B and C, and scenario 2, ghost introgression (dotted orange arrow) between ghost lineage X and species B. Bottom panel. Left: the three possible topologies representing the relationship between species A, B and C. Right: The mean divergence times T1 and T2 were computed for both scenarios and for all genes supporting either the species tree topology ((A,B),C) (black rhombus) or the two discordant topologies ((B,C),A) and ((A,C),B) (green rhombus; whiskers standard error of the mean, *** for t-test with P-value < 2.2e-16). Data underlying this figure can be found on Zenodo (doi: 10.5281/zenodo.6901799) and GitHub (https://github.com/theotricou/Ghost_branch_length/tree/main/2_Anopheles).

We observed that under scenario 1, divergence times in gene trees with discordant topologies were on average shorter than in gene trees with the same topology as the species tree (Fig 4). This is in agreement with the expectations of the *Anopheles* study (citation above). However, when introgression came from a ghost lineage from outside the ingroup (such as in scenario 2), we observed the opposite result, with gene trees with discordant topologies exhibiting longer divergence times than gene trees with the same topology as the species tree. In this case, it is incorrect to consider that the true species tree topology is the one supported by the genes with longest divergence times.

To complete this analysis, we performed another type of simulation where no alternative scenario was imposed (unlike scenario 2 above). This addresses the plausibility of scenario 1 versus scenario 2 in the presence of ghost lineages. We simulated the evolution of genomes (with random interspecies introgressions) along a randomly generated species tree containing ghost lineages (extinct species in this simulation). We obtained gene trees and we computed the divergence times (T1 and T2) of the genes having various topologies for 3 clades considered (A, B, C) (see S1 Material). We did the same simulation of genome evolution on a reduced species tree obtained by removing extinct (ghost) lineages from the original one. We observed, in accordance to our previous results, that under this setting, when simulations were performed on a species tree containing ghosts, divergence times (T1 and T2) in genes that were not following the species tree topology were higher than those of genes following the species topology (S1Fig). Again, the expectation that “gene trees constructed from non introgressed sequences [would] show deeper divergences than those constructed from introgressed sequences” [13] was violated. Back to Fontaine et al. (2015) this suggests that the “true” branching order, as the authors call the searched species tree, could be the one with the shortest tree height, compatible with autosomal regions being in accordance with the species tree topology. Confirming or disconfirming such an alternative scenario is out of the scope of this work. It would require a re-analysis of a substantial set of arguments. What we point out is a weakness of the phylogenetic arguments presented above, which are important ones in the construction of the proof. As far as we know, no study on this group specifically tested the hypothesis of a ghost introgression.

### Using branch lengths to order acquisition events (the stem-length method) is misleading in the presence of ghost lineages

Recent studies have used the so-called “stem-length method” to reconstruct the relative timing of acquisition of different genes by a proto-eukaryotic cell during eukaryogenesis [14,16]. This method was also used to characterize the relative timing of acquisition of genes involved in the elaboration of the chloroplast membrane [20]. The stem-length method, shown in Fig 5, relies on the expectation that early acquisitions of genes by a given recipient lineage should result in longer branches (or stems) at the base of this lineage in the corresponding gene trees than late acquisitions (Fig 5, top). This approach was used to address the long-standing question of the early or late acquisition of mitochondria during eukaryogenesis [14]. The authors of this study concluded that the shorter stems in the trees of eukaryotic genes of alphaproteobacterial origin, as compared to genes of other bacterial origins associated with other components of the eukaryotic cell, supported a late acquisition of mitochondria. Precisions [31] and critics [32] about some of the theoretical, methodological and statistical aspects of this study have already been formulated, and commented on by the authors [33]. Here we focus on an overlooked one: ghost lineages. Their existence was mentioned earlier as a possible “caveat[s] accompanying conclusions coming from stem-length methods” [31], but their impact on the method was never evaluated.

**Fig 5.**
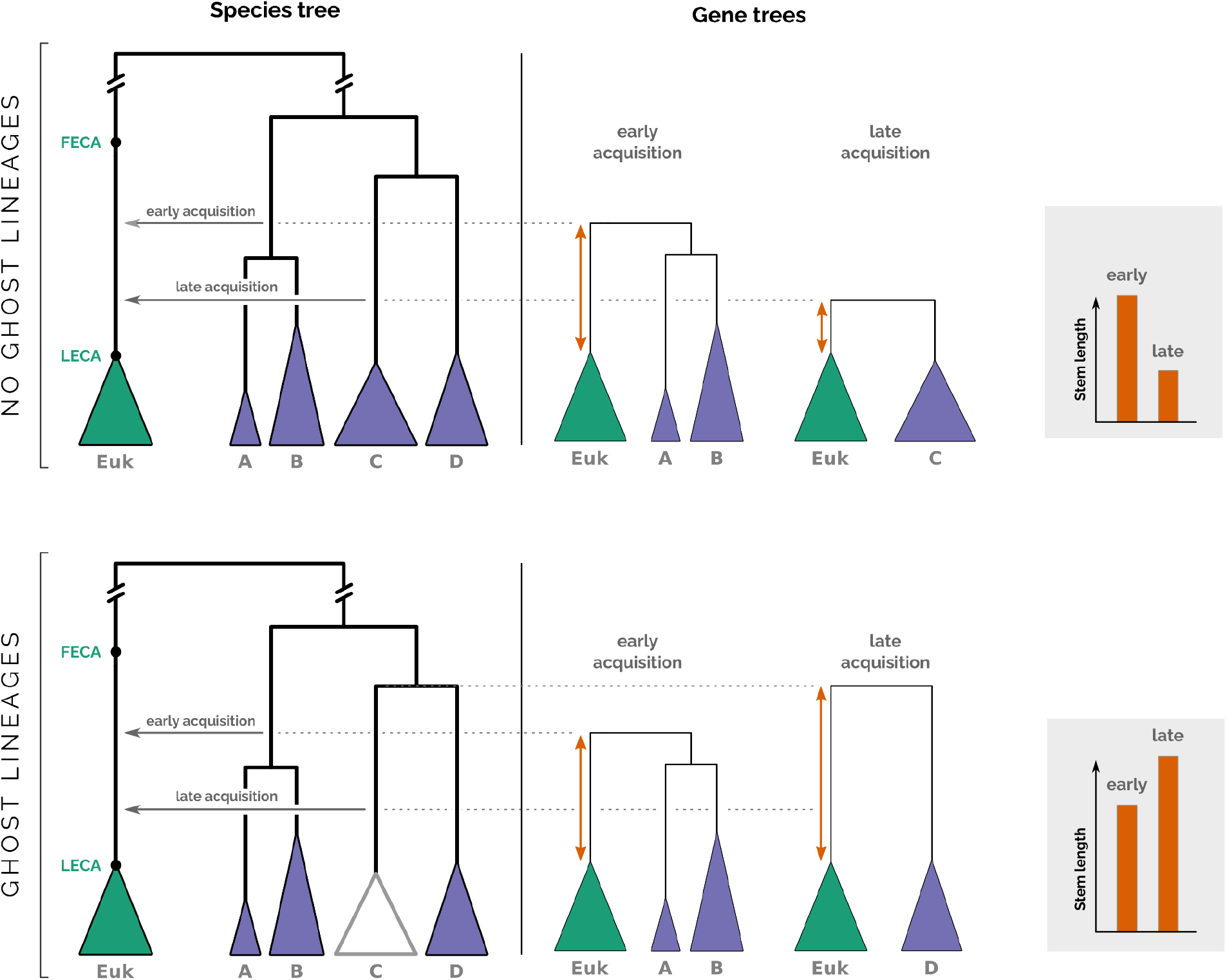
Illustration of the impact of ghost lineages on the relative timing of genomic acquisitions with the stem-length method. Under the hypothesis of all clades being sampled, i.e. there are no ghost lineages (top), an early acquisition produces a gene tree with a long stem, while a late acquisition produces a gene tree with a short stem (orange arrows show stem lengths, shown in the grey box on the right). If a clade is missing, i.e. there are ghost lineages (bottom), the opposite observation can be made. Because the donor lineage (here the ancestor of C) left no descendants, and because it split from the rest of the clade (D) before the time of the early acquisition event, the correlation between stem lengths and acquisition time is reversed: early acquisition leads to a shorter stem length than late acquisition (grey box).

If ghost lineages are taken into account, the expectations of the stem-length method can be totally reversed. If the donor lineage has no descendants, either because they all went extinct, because they have not been discovered, or because they were not considered in the study, then stem lengths will be determined not by the time when the transfer occurred but by the divergence time between the missing (ghost) clade from which the transfer originated and its closest sampled relative (Fig 5, bottom). Under these circumstances, the correlation between order of acquisition and stem-length is lost. To quantify this effect and have an insight into the validity of the results using the stem length method, we performed a simulation where species trees with 1,000 leaves were generated following a birth-death process and pairs of acquisition events were repeatedly sampled in the tree to mimic early and late acquisitions. We then sampled a proportion of the extant species (between 1% and 10% of the total) and looked at the effect of sampling on the predictions of the stem-length method regarding the order of events (Fig 6). We observed that when 10% of species were sampled, ∼33.5% of the predictions were wrong (predicting that event A occurred before event B when the opposite actually happened). This proportion reached almost 50% (the maximum possible, equivalent to a random prediction) when 1% of the species were sampled (Fig 6A).

**Fig 6.**
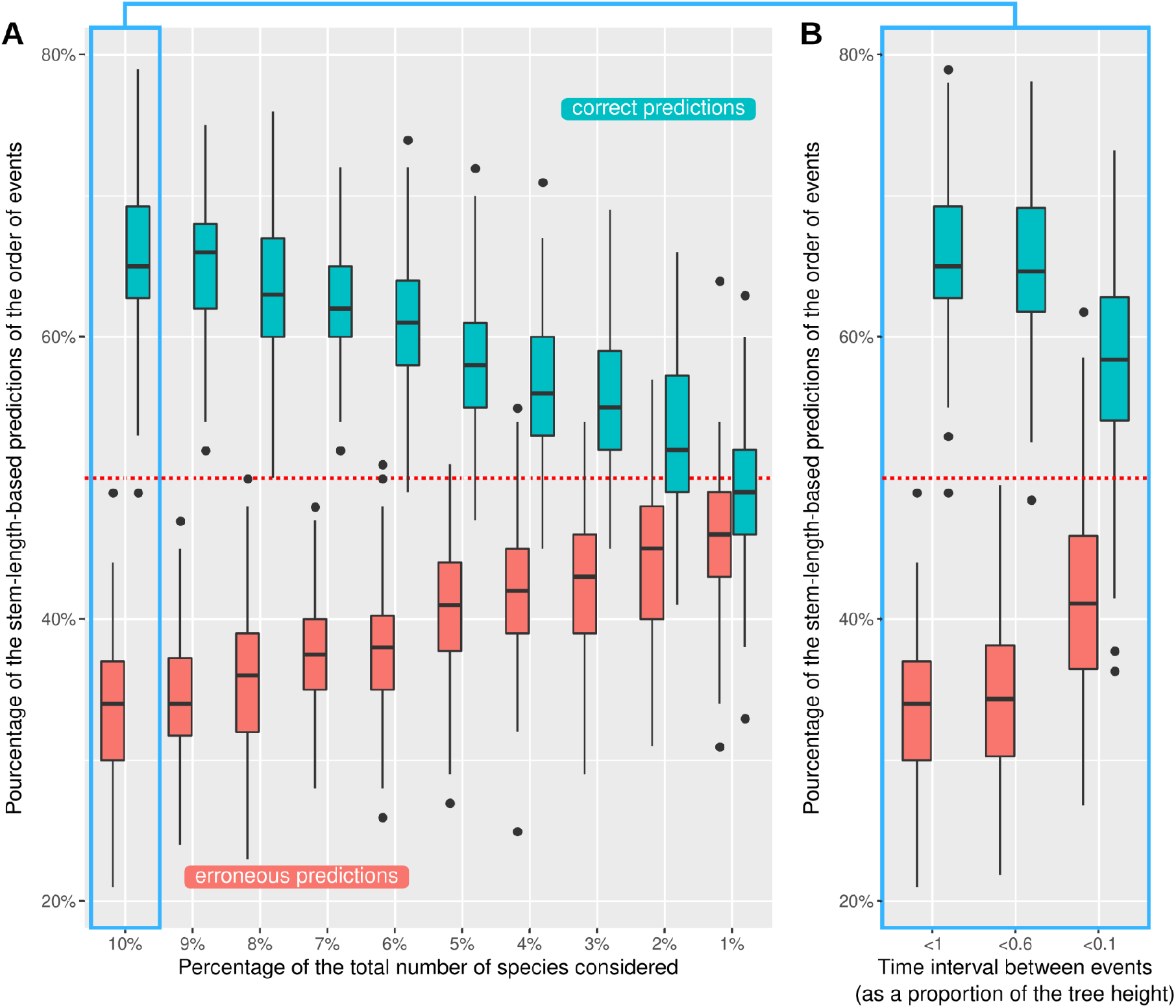
Impact of sampling on the stem-length method. Effect of the proportion of species sampled (**A**) and the length of the time interval between acquisition events (**B**) on the percentage of right (blue bars) and wrong (red bars) predictions of the order of events using the stem-length method. The dashed red line represents what would be observed if predictions were random. Cases where the stem-length method could not order events (because stems had the same length) were removed, which explains why the blue and red bars do not add up to 100%. Data underlying this figure can be found on Zenodo (doi: 10.5281/zenodo.6901799) and GitHub (https://github.com/theotricou/Ghost_branch_length/tree/main/3_Stem-length).

We also observed that the error increased when the time interval between the two events was shorter. Indeed, nearly 41% of predictions were wrong when the time interval between events was less than 10% of the tree height with 10% of species sampled (Fig 6B).

## Discussion

Branch-length-based approaches are versatile methods for studying different aspects of horizontal gene flow (HGF) at different scales, from intraspecific introgression (between populations) to transphylum gene transfers. These approaches rely on the expectation that any HGF should result in shorter (phylo)genetic distances between donor and recipient in trees of the transferred genomic sequence(s) compared to trees of other (presumably vertically transferred) sequences from the same taxa. However, this only holds if all lineages, or at least the sister lineages of all lineages under consideration, are present in the study. Indeed, as demonstrated here, HGF can result in longer (phylo)genetic distances in trees of the transferred genomic sequence(s) if the donor is absent. Therefore, it appears crucial to take ghost lineages into account when interpreting the result of branch-length-based HGF inference methods.

The simulations performed here are simple, but they illustrate the impact that ghost lineages can have on approaches as diverse as the D_3_ test [9], the test developed for finding the correct branching order in species groups [13] and the stem-length method for ordering events [14]. First, our simulations demonstrate that under the realistic assumption that there are many ghost lineages, not only does the likelihood that HGF comes from unknown species increase, but also the possibility that the identification of both the donor and the recipient of the transfer is erroneous, leading to the identification of two lineages that have nothing to do with this process. Thus, when interpreting the results of introgression tests like D_3_ [9], the possibility of introgression from ghost lineages from outside the three taxa considered should be systematically taken into consideration as a possible alternative scenario and should be considered to be (at least) as probable as the usual interpretation.

Second, we show that HGF from a ghost lineage could in some cases increase branch lengths in a gene tree, instead of decreasing them as is commonly expected. This absence of an unambiguous pattern suggests that branch lengths may not be suitable for identifying appropriate markers for phylogenetic reconstruction as was done in Fontaine *et al*. [13]. Introgressions are frequent in the *Anopheles* genera, so that the possibility of an introgression from an unsampled (known, unknown or extinct) species does not seem unlikely.

The article by Fontaine et al [13] is part of an important literature supporting a massive autosomal introgression in the *Anopheles gambiae* complex. Other analyses performed on the same set of species using different methods postulated autosomal introgressions involving *An. arabiensis, An. gambiae* and *An. coluzzii* [34–38]. However, none of these studies considered the possible role of ghost lineages in giving the observed patterns of introgression. Just like we demonstrated with the reanalysis of Fontaine et al. [13] that an alternative conclusion was possible, re-analysis of these studies with considering the possible impact of ghosts can change the interpretation of their results as well. For instance in Thaworwattana et al. [38] the introgression detection method is based on divergence time, which we demonstrate to be highly sensitive to the influence of ghost lineages, and the arguments used to support the species tree topology agreeing with the X chromosome seem compatible with the opposite hypothesis (species tree topology agreeing with the autosomes) if ghosts are considered.

Third, because HGF can increase or decrease branch lengths depending on the sampling effort and the proportion of ghost lineages, we show that the stem-length method that infers the relative timing of transfer events is prone to errors. To quantify this effect on biological data, it is necessary to answer the following questions regarding any clade where the stem-length method is to be applied: 1) How likely is it that the lineages from which the transfers originated have no descendants?, and 2) How far apart are the events under investigation, relative to the total timespan considered?

We can provide the following elements concerning the biological data for which the stem-length method was devised and on which it was applied originally [14,16]. The acquisitions in question are genes of bacterial origin transferred to protoeukaryotes after the appearance of the First Eukaryotic Common Ancestor (FECA) and before that of the Last Eukaryotic Common Ancestor (LECA), *i*.*e*. between ca. 2.3-2.7 and ca. 1.1-1.2 BYA (reviewed in Chernikova et al. 2011). It is difficult to infer the macroevolutionary history of bacteria, especially at such deep evolutionary timescales, but it is clear that most lineages living during that period went extinct [40] and some went extinct during mass extinctions [41]. In addition, it is clear that most extant bacterial lineages are still unknown [19,40] and may not be scattered uniformly across the known diversity. Indeed, unsampled lineages are likely to form major clades, as illustrated by the discovery of a complete new phylum (CPR) in 2016 [42].

Given that only a small fraction of the diversity (extinct and extant) is known, we may consider the stem-length method as equivalent to a random or arbitrary choice of the relative timing of acquisition events.

In this work, we explored how ghost lineages can affect methods based on branch-lengths to study various evolutionary events in the presence of HGF. The confounding effect that ghosts can have on these methods has been acknowledged earlier [9,31]. It has, however, always been implicitly considered less likely. We showed that the main interpretation of such methods should consider ghosts as the primary actors, and that doing so radically changes the interpretations of the tests and therefore the main conclusions of the studies on which it relies.

It is also important to mention that alternative methods have been developed in recent years to be able to explicitly take ghost lineages into account when predicting introgression scenarios, and to ultimately allow predicting the existence of ghost (archaic) species themselves [43,44]. Of note, the use of these methods is often restricted to datasets for which the availability of archaic genomes can help validate the predictions made, such as primate [including humans, 45,46] or bear [47].

One direction of research could be the development of methods to detect ghost lineages or to disentangle ghost introgression from known lineage introgression. This would increase our knowledge of ghost lineages which may help to solve the interpretation issues we just demonstrated. However we feel that it is not a necessity to “unghost” the lineages to handle them in the interpretations of current methods: given their overwhelming importance, it is more promising to highlight the fact that most HGF-detection results are interpreted without considering ghost lineages, and that failure to consider their impact on the interpretation of such tests impair their usefulness and lead to erroneous conclusions. As such, and considering the huge number of invisible lineages in real life, we suggest that the results of gene flow-related methods should be interpreted with the signature of ghost lineages as the foremost hypothesis.

## Material and Methods

### The effect of ghost lineages on the D_3_ test for introgression

#### Simulating species trees and introgressions

200 random species trees with a birth-death model were simulated using *Zombi* [48]. Speciation and extinction rates were fixed at 1 and 0.9, respectively. Simulations were run until 40 extant species were reached then 20 species were sampled. We converted the topology of the species tree into a suitable format for the coalescent simulator *ms* [49] with a custom python script: branch lengths were converted into units of generation and the age of the root of the trees was fixed at 10^6^ generations. To simulate introgression, a single migration event was imposed over one generation for a fraction **f** = 50% of the donor population targeting the recipient population. This migration rate was used to ensure that introgression detection using D_3_ would not be biased by false positives. Then, for each species tree, we used *ms* implemented in the R package *Coala* <u>[50]</u> to simulate 1,000 gene trees evolving in populations of fixed size (Ne) of 100,000 individuals.

#### Computing the D_3_ statistic

To compute the D_3_ statistic for three lineages with the species tree topology ((P1,P2),P3), we calculated D_13_ and D_23_, the sum of the distance (or branch length) separating P1 and P3 and P2 and P3, respectively, across all gene trees (1,000 in total). We then computed the D_3_ statistic using equation (1). This was done for each trio of lineages ((P1,P2),P3) in each species tree simulated. D_3_ significance was tested by bootstrap resampling of 1,000 gene trees with 1,000 replicates. We then calculated the Z-score and considered that D_3_ was significant if Z > 3 or Z < 3, following Green et al. (2010). Finally, as we tracked the true donor and recipient lineages in each simulation, we assessed for each D_3_ whether the test was significant due to an ingroup introgression or a ghost introgression event from outside the three species considered.

### Using branch lengths to determine the correct species tree topology

#### Species tree simulation with ingroup or ghost introgression

A species tree was simulated using *Zombi* [48]. Unless otherwise stated, parameters were the same as the one used in the previous section. The simulation was run until 16 extant species were reached. On this tree, three species with the topology ((A,B),C) were arbitrarily chosen. All others were considered to be ghost lineages. We then converted the topology of the tree into an *ms* readable tree for coalescent simulation. Two datasets were generated with *ms*; in both datasets the number of generations separating the tip from the root of the tree was fixed to 5×10^6^ and a migration event was imposed for a fraction ***f*** = 20%. For the first dataset, migration took place from B to C, *i*.*e*. between two extant species (ingroup introgression). For the second dataset, migration took place between a randomly sampled ghost lineage outside of the triplet phylogeny and B (ghost introgression). For both models, we simulated 1,000 gene trees evolving in populations of fixed size (Ne) of 100,000 individuals.

#### Branch length in gene trees

We identified the gene trees that were congruent with the ((A,B),C) topology (corresponding to the branching order of the species tree) and those that corresponded to one of the two discordant topologies ((B,C),A) or ((A,C),B) that can arise from ILS or introgression. Subsequently, for both introgression models described above and for each gene tree, we computed the value of species divergence times T1 and T2 (see Fig 4) following the equation from Fontaine et al. (2015) (see supplementary material S3.2 in Fontaine et al. (2015)).

### The effect of ghost lineages on the inference of the relative timing of gene acquisitions using the stem-length method

The order of acquisition events from simulated trees with and without ghost lineages was compared to the predicted order of events using the stem-length method [14,16]. Simulations were carried out as follows:

- 100 trees under a birth-death process (speciation rate = 1, extinction rate = 0.5) were generated using the *rphylo* function in the R package *ape* [52]; simulations were run until trees reached 1,000 leaves.
- Two points were randomly sampled from each tree representing two origins of transfer (or acquisitions) towards the same recipient lineage; we recorded their timing and the time interval (*dt)* between them (as a fraction of the total tree height).
- Trees were pruned by sampling a proportion *p* of the leaves from each tree.
- We evaluated the new order of events on the pruned tree using the stem-length method.
- We recorded whether the order of events before and after pruning was the same (1), was different (0) or could not be determined (*i*.*e*. events occurred at the same time) (NA). The latter occurs if the two events lead to equal stem lengths after sampling.
- The proportion of each (1, 0, NA) out of the 100 replicates was calculated.

The proportion of sampled leaves *p* ranged between 1% (10 leaves) and 10% (100 leaves), in 1% increments. We performed 100 replicates to obtain a variance for the observed proportions of correct and erroneous predictions of the order of events.

To explore the effect of the time interval (*dt*) between events on the proportion of erroneous predictions, we subsampled the pairs of events with *dt* < 1, *dt* < 0.6 and *dt* < 0.1 and recomputed the proportion of correct and erroneous predictions each time. This analysis was carried out with *p*=10%.

## Acknowledgments

We thank Adrián A. Davín for sharing his insight on performing simulations and Gergely J. Szöllősi for useful discussions. This work was supported by the French National Research Agency (Grants ANR-18-CE02-0007-01 and ANR-19-CE45-0010). Simulations were performed using the computing facilities of the CC LBBE/PRABI.

## Software Availability

All codes used to generate and analyze the simulations performed in this study are available on Zenodo (doi: 10.5281/zenodo.6901799) and GitHub (https://github.com/theotricou/Ghost_branch_length.git).

## Supporting information

### S1 Material. Additional simulations for exploring the effect of ghosts on the method employed by Fontaine et al. (2015) to find the correct branching order of *Anopheles* species

To explore the effect of ghost lineages on the observed divergence times in gene trees following or not the species tree topology, but without choosing arbitrarily a specific scenario as Fig 4 (main text), we used the approach detailed hereafter, using the software Zombi (Davín et al. 2020):

- We simulated a species tree containing extinct (ghost) species. This tree was obtained using the **Tp** mode of Zombi, with a speciation and extinction rates of 0.15 and 0.09 respectively. The lineage number was set to 20 species for the 140 first units of times then to 3 species for the last 10 units with a turnover set to 0.05. This produced a complete tree of 119 species of which only 10 were extant (S1B Fig). These extant species were grouped arbitrarily in three clades, A, B and C, with topology ((A,B),C).
- We produced a new version of this species tree that did not contain ghosts by simply pruning them out (S1A Fig).
- On each version of the tree, we simulated the evolution of a genome evolving along its branches, letting interspecies introgression occur. Genomes with an initial size of 3000 genes were made to evolve in both trees using the **G** mode in Zombi. Introgression was simulated by setting the transfer rate of genes to 5 for the tree with extinct species and 10 for the other one. The probability of replacement transfers was set to 1. The extension parameter of the transfer events was set to 1, meaning that every time that a transfer event occured it affected only a single gene.
- We recovered the gene trees at the end of the simulations and we grouped them according to which topologies they supported: either the topology reflecting the species history: ((A,B),C) or one of the two topologies resulting from introgressions, ((A,C),B) and ((B,C),A). For each of the three topologies and for each simulation, mean T1 and T2 were computed
- The results (S1 Fig) were presented following Fig 3 (A and C) of (Fontaine et al. 2015).

**S1 Fig.**
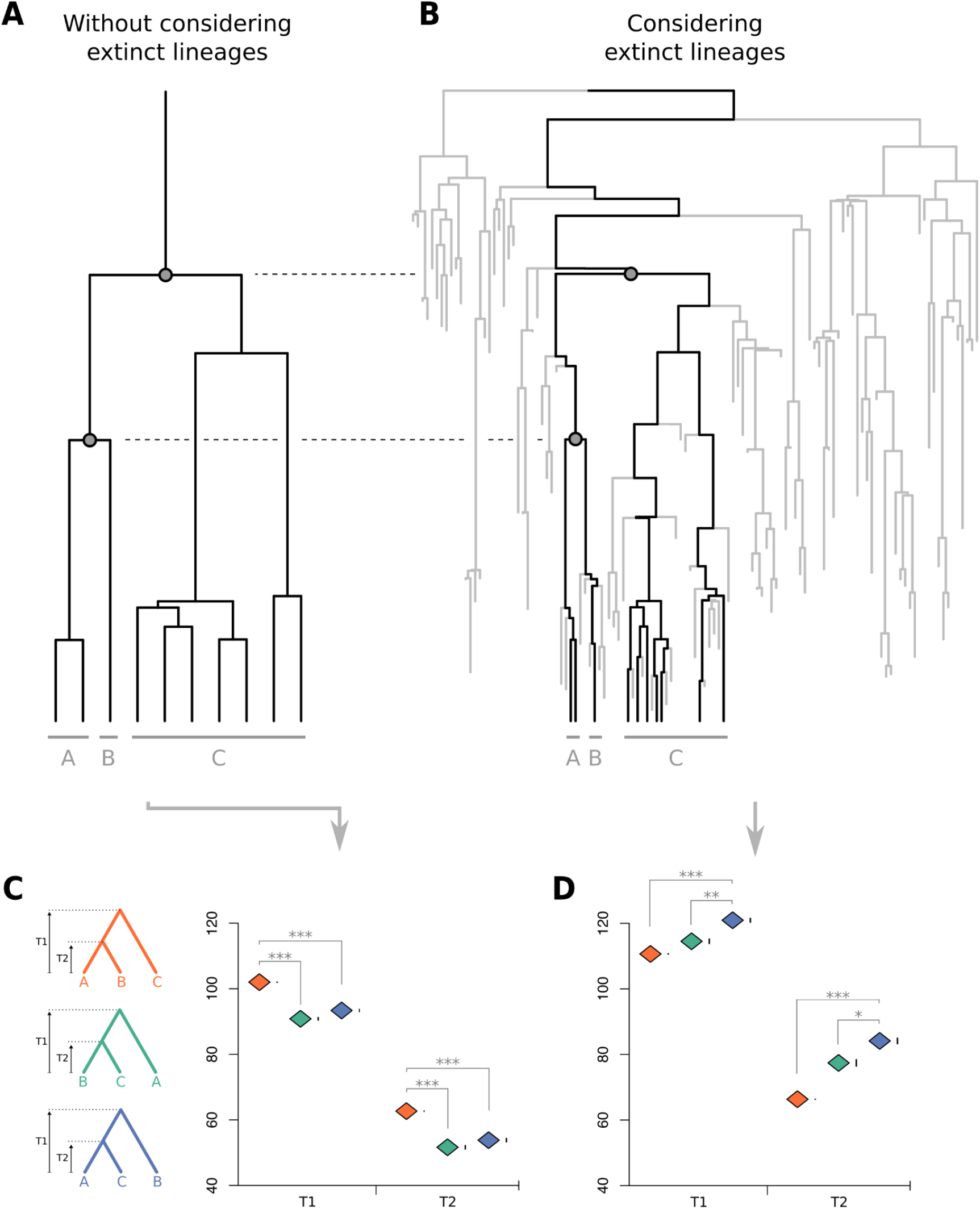
Effect of ghost lineages on observed divergence times in gene trees following or not the species tree topology. The evolution of genomes (3000 genes) was simulated on two trees, one with only extant species (**A**), the other with all species (**B**). Mean T1 and T2 were computed for all genes supporting each one of the three possible topologies (orange, green and blue trees).The true topology (orange) is the one supported by the trees with the highest mean T1 and T2 when considering only extant species (**C**), but when considering also extinct lineages (**D**), the gene trees with the highest T1 and T2 support instead an incorrect topology. One-tailed t-test p-values: * < 0.01; ** < 0.001; *** < 0.0001. Vertical bars on the right of diamonds represent standard error. Data underlying this figure can be found on Zenodo (doi: 10.5281/zenodo.6901799) and GitHub (https://github.com/theotricou/Ghost_branch_length/tree/main/2_Anopheles/Sup_dataset).

